## Supplementary material for "A novel human pluripotent stem cell-based gene activation system identifies IGFBP2 as a mediator in the production of hematopoietic progenitors in vitro"

### Supplementary Figures

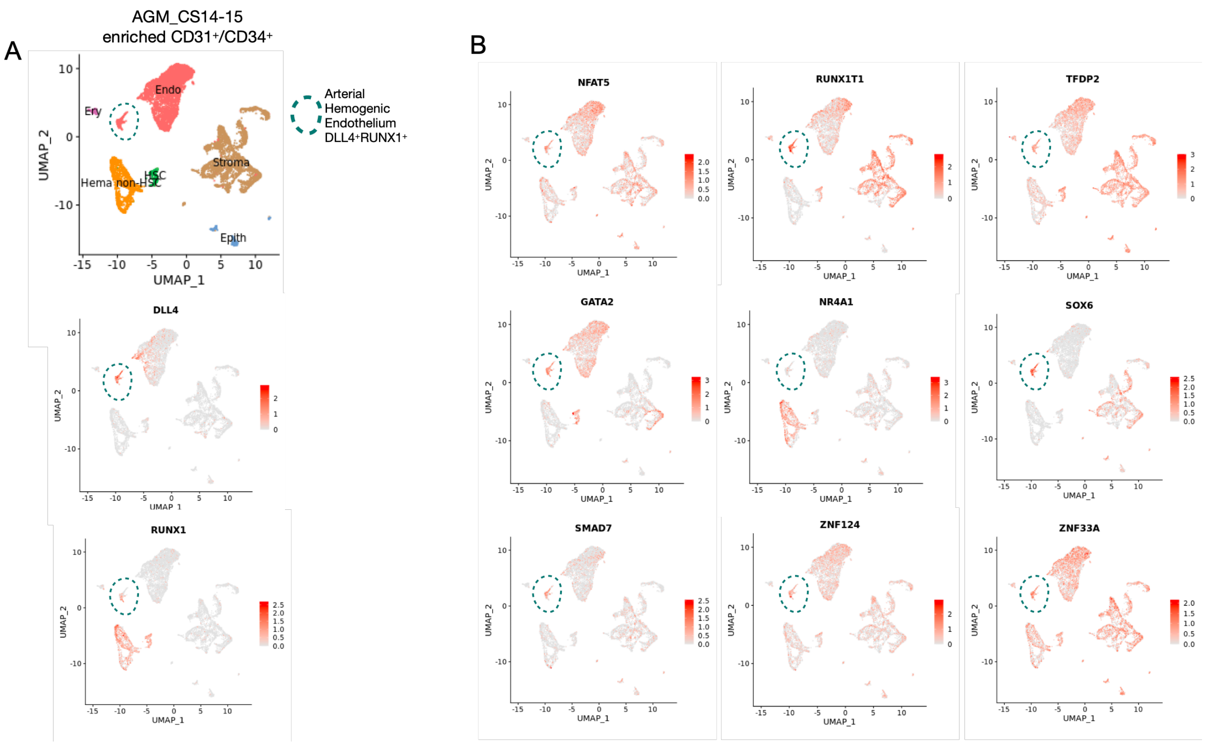


### Supplementary Figure 1

**A** – Single-cell transcriptomic analysis of developing AGM collected from human embryos at Carnegie Stages14 and 15 enriched for CD31$^{+}$ and CD34$^{+}$ cells from Calvanese et al 2022, Nature. Arterial hemogenic endothelium, dashed line, is identified by the colocalization of *DLL4* and *RUNX1*. **B** – Gene expression analysis of the 9 selected target genes in the developing human aorta, showing their expression in the hemogenic endothelium.

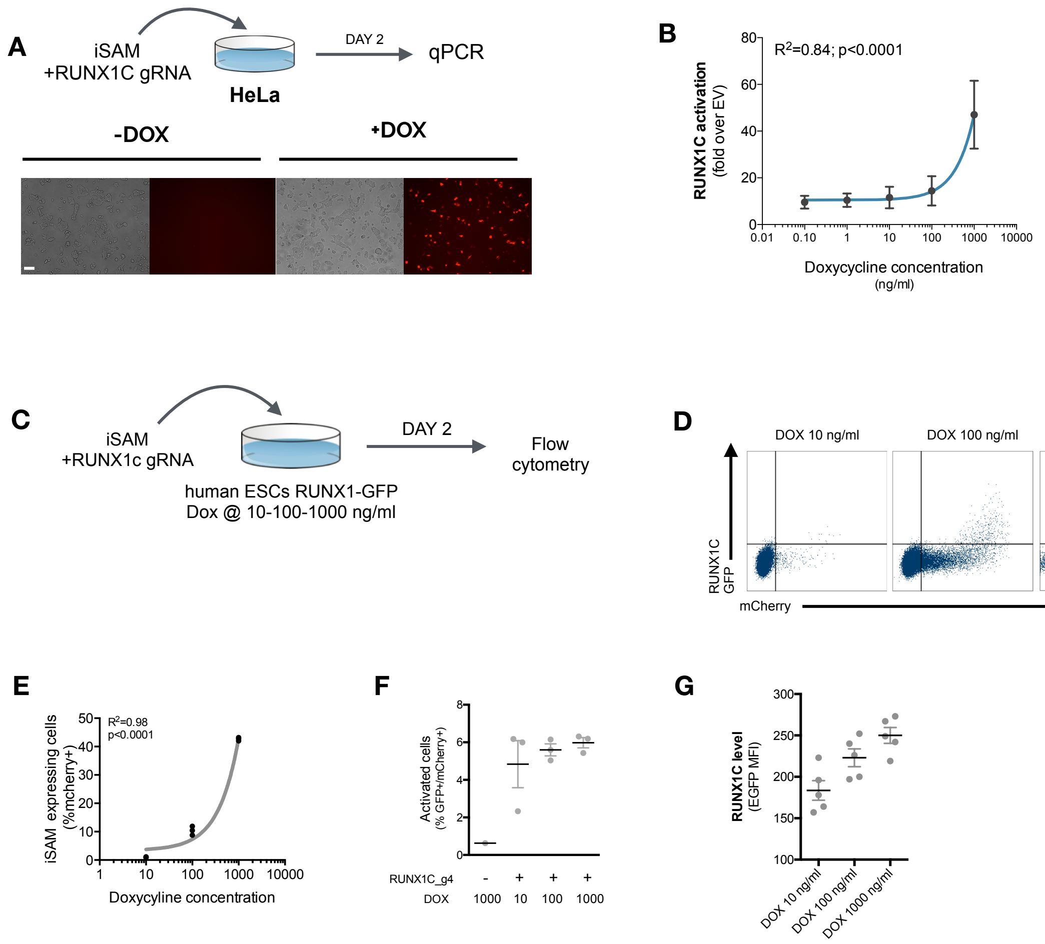


**Supplementary Figure 2**

**A** – Schematic of the iSAM mediated activation of RUNX1C by transient transfection in HeLa cells with the iSAM vector and the RUNX1C gRNA; fluorescent microscopy demonstrating the expression of the mCherry tag. **B** – Linear regression of *RUNX1C* RNA expression in relation to the concentration of DOX added to HeLa cells (n=3). **C** - Schematic of the iSAM mediated activation of the hESCs RUNX1C-GFP reporter cell line by transient transfection of the iSAM vector and RUNX1C gRNA. **D** – Flow cytometry analysis of RUNX1C-GFP expression and mCherry in the hESCs RUNX1C-GFP reporter line, after exposure to different DOX concentration. **E** - Linear regression of the mCherry tag expression in relation to the concentration of DOX added (in ng/ml) to the hESCs RUN1C-GFP reporter line (n=3). **F** – Percentage of activated cells (GFP$^+$mCherry$^+$) in presence of different concentration of DOX (in ng/ml) (n=3). **G** – RUNX1C single cell expression level analysed by flow cytometry in hESCs exposed to different DOX concentration (n=4).

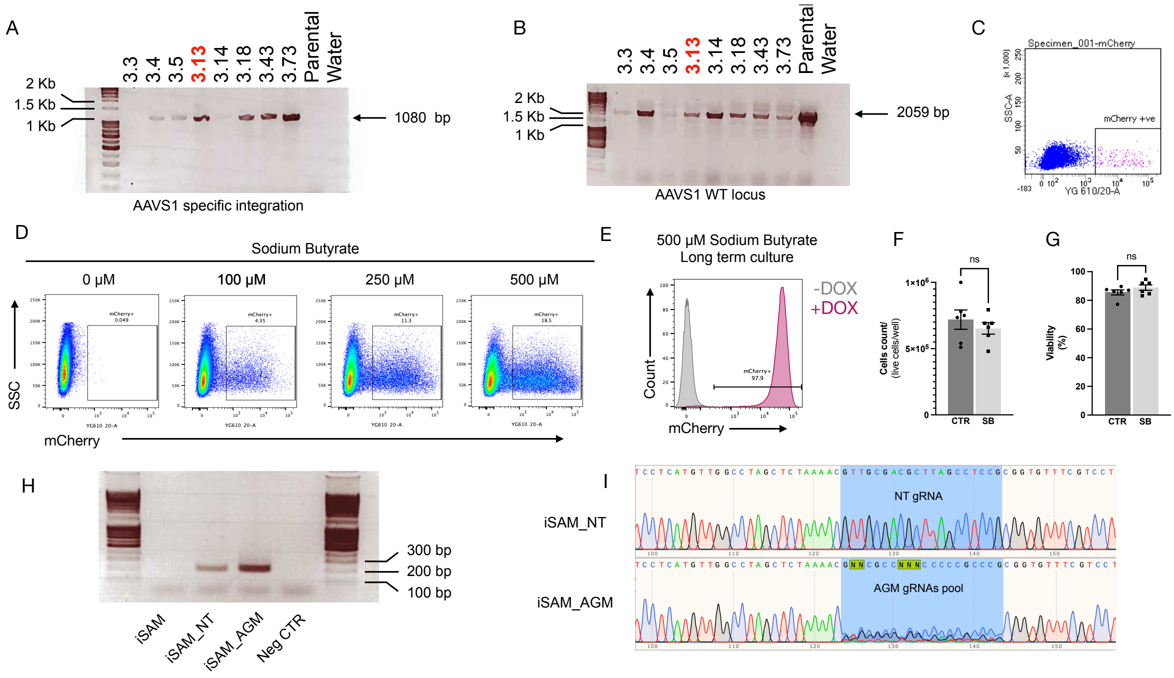


### Supplementary Figure 3

**A** – PCR screening of the clones obtained from the AAVS1 targeting showing specific integration with amplification across the 5' end, and **B** - the WT locus. Clone 3.13 was selected for the study and referred to as iSAM. **C** – Flow cytometry analysis of mCherry+ cells upon DOX addition in hiPSCs with iSAM targeted into the AAVS1 locus following maintenance. **D** – Flow cytometry analysis of the expression of the mCherry tag, in hiPSCs with iSAM targeted into the AAVS1 locus, following 48h Sodium Butyrate and DOX treatment at different concentrations. **E** - Flow cytometry analysis of the expression of the mCherry tag upon DOX addition, in hiPSCs with iSAM targeted into the AAVS1 locus, maintained in the presence of 500 µM Sodium Butyrate. **F** – The cell count of iPSCs maintained in the presence of Sodium Butyrate shows no significant differences (Mann-Whitney unpaired T-test, p=0.45). G – Cell viability was comparable when iPSCs were maintained in control conditions or in the presence of Sodium Butyrate. (Mann-Whitney unpaired T-test, p=0.46) **H –** PCR screening for the integration of the gRNA into the genome of the iSAM hiPSCs line (from left to right: iSAM line before infection, after infection with non-targeting gRNA, after infection with AGM library and water negative control). **I –** Sanger sequencing trace of the amplicons obtained from the iSAM line infected with the non-targeting gRNA (top) or the AGM library (bottom).

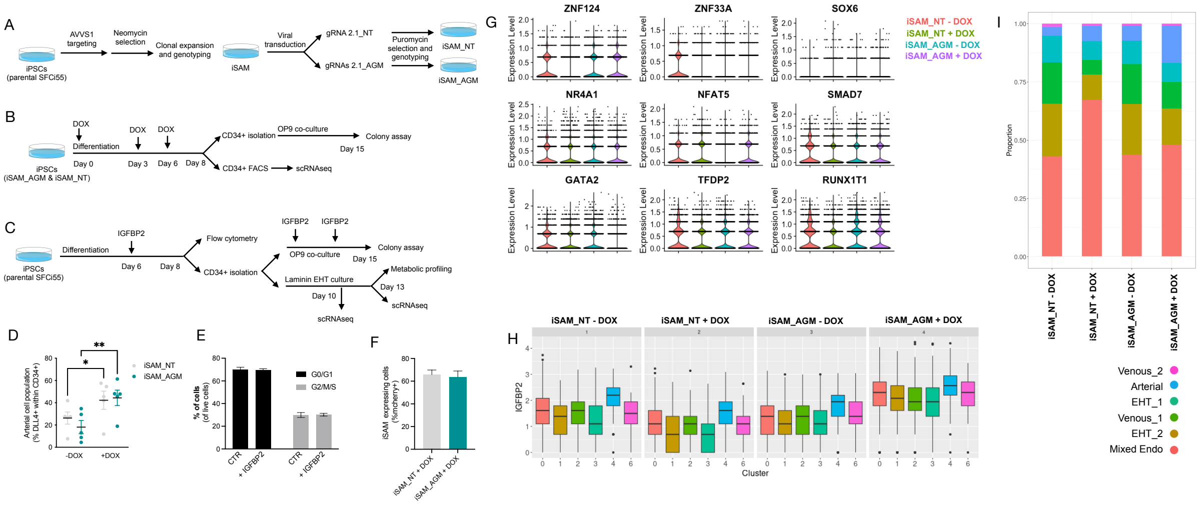


### Supplementary Figure 4

**A** – Schematic of the derivation of the iSAM cell line by ZNFs mediated targeting of the AAVS1 locus and the subsequent derivation of the iSAM_NT (containing the non-targeting control gRNA) and iSAM_AGM (containing the gRNAs for the target genes) by viral transduction of the gRNAs. **B** - Schematic of the differentiation protocol of the with activation of the target genes, used for both the control line iSAM_NT and iSAM_AGM. **C** - Schematic of the IGFBP2 functional validation experiment. **D** – Expansion of the arterial population marker by membrane expression of DLL4+ following targets' activation, quantified by flow cytometry at day 8 of differentiation (* p = 0.0190, ** p = 0.0011, Sidak's Two-way ANOVA). **E** - Flow cytometry analysis of the cell cycle analysis of suspension progenitor cells was obtained following the OP9 coculture of cells treated with IGFBP2 and control at day 13. **F** - iSAM expressing cells at day 8 of differentiation upon DOX treatment. **G** - Target genes' expression profile across the different libraries, the colour legend for the libraries is on the top right. **H** - IGFBP2 expression profile across the cell clusters and in the cell lines and conditions indicated in the title (cluster legend same as in I). **I** - Contribution of each of the libraries to the cell clusters, grouped by libraries (cluster legend at the bottom).

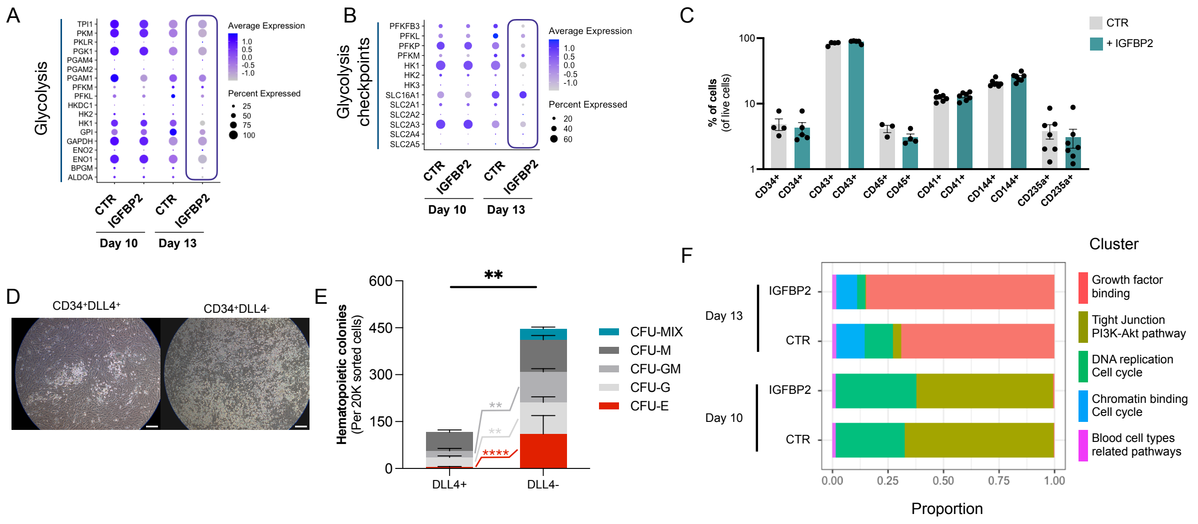

**Supplementary Figure 5**

**A** – Dot plot showing the expression profile of the genes coding for the enzymes mediating glycolysis in CTR and IGFBP2-treated cells at day 10 and 13. **B -** Dot plot showing the expression profile of the genes coding for the checkpoints of the glycolysis in CTR and IGFBP2-treated cells at day 10 and 13. **C -** Percentage of cells expressing different markers following OP9 coculture of CD34+ cells (ns for all markers, Kruskall-Wallis One-way Anova). **D -** OP9 co-cultures of CD34+DLL4+ and CD34+DLLL4- (scale bar indicates 50 µm), and in **E** their colony formation capacities following one week of OP9 co-culture (Mixed-effect analysis, Sidak's post-test, **** p<0.0001, ** p<0.01). **F -** Cluster composition in cells treated with IGFBP2 data shows the composition at day 10 and day 13.
